# Mechanistic Modeling of Sleep-Wake Transitions via Circadian-Modulated Threshold Dynamics

**DOI:** 10.1101/2025.07.10.664059

**Authors:** Chenggui Yao, Dongping Yang

## Abstract

Human sleep-wake cycles emerge from complex interactions between homeostatic sleep pressure and circadian rhythms. In this study, we extend the Phillips-Robinson model by introducing circadian-dependent dynamic thresholds for sleep and wake transitions, yielding a more physiologically grounded framework for sleep regulation. Using bifurcation analysis, we show that the transition from sustained wakefulness to rhythmic sleep-wake cycles is governed by a saddle-node on invariant circle (SNIC) bifurcation, and that these oscillations become entrained to external 24 h light-dark cues. We analytically derive circadian-modulated sleep and wake thresholds, revealing how the interaction between circadian and homeostatic drives governs sleep-wake transitions. Our model captures key physiological phenomena, including: (1) the onset and entrainment of sleep-wake rhythms, (2) immediate sleep onset and partial rebound following sleep deprivation, and (3) sleep fragmentation under shift work-like conditions. These results offer new mechanistic insights into how circadian misalignment alters sleep timing and quality. Together, our findings establish an updated theoretical framework for modeling sleep-wake regulation in both natural and disrupted environments, with implications for shift work management, sleep disorder interventions, and personalized chronotherapy.

**Author summary:** Sleep and wakefulness are regulated by a complex interaction between circadian rhythms and homeostatic sleep pressure. While existing computational models have provided valuable insights, most rely on fixed or heuristic thresholds for sleep and wake transitions that lack direct physiological basis, limiting their ability to account for the full range of sleep behaviors–especially under disrupted conditions. In this study, we substantially update the computational framework in which sleep-wake transitions are governed by dynamically circadian-dependent thresholds. It captures how internal biological time shapes sleep patterns and explains key phenomena such as sleep rebound following deprivation and sleep fragmentation during shift work. It provides new insights into how circadian misalignment leads to sleep disorders and lays the groundwork for more effective, personalized interventions.

## I. INTRODUCTION

Sleep and wakefulness are fundamental behavioral states regulated by complex physiological mechanisms. Their coordination is governed primarily by two interacting processes: the circadian rhythm, which operates on an approximately 24-hour cycle, and the homeostatic sleep pressure, which accumulates during wakefulness and dissipates during sleep. Borbély’s seminal two-process model conceptualized this interplay, proposing that the timing and duration of sleep and wake episodes are determined by the interaction between circadian and homeostatic drives, mediated through circadian-dependent thresholds for sleep and wakefulness [1]. However, in this framework, the sleep and wake thresholds were imposed in an ad hoc manner, lacking direct physiological justification [2].

Recent studies have proposed several physiological mechanisms that support the existence of circadian-dependent thresholds. The suprachiasmatic nucleus (SCN), the brain’s central circadian pacemaker, exerts time-of-day-specific influence on sleep- and arousal-regulating regions such as the ventrolateral preoptic area (VLPO) [3]. This modulation is coordinated through rhythmic hormonal outputs such as melatonin and cortisol, which shift neural excitability and alter the propensity for sleep or wakefulness [4]. Moreover, circadian control of thermoregulation–including core body temperature fluctuations–further influences sleep onset and maintenance [5]. Notably, circadian phase gates the responsiveness of the sleep-wake system to homeostatic sleep pressure (e.g., adenosine), such that an identical level of sleep pressure may yield different behavioral outcomes depending on internal time [6]. At the molecular level, clock gene expression in extra-SCN sleep-regulatory nuclei suggests the presence of local circadian oscillators capable of dynamically modulating neuronal excitability and network dynamics [7]. Collectively, these findings provide a physiological foundation for modeling sleep and wake thresholds as circadian-dependent, rather than arbitrary, constructs.

The Phillips-Robinson (PR) model offers a more physiologically grounded approach by explicitly defining two biologically meaningful thresholds that govern transitions between sleep and wake states [8, 9]. These thresholds have proven essential for accurately capturing sleep-wake dynamics and have enabled a range of practical applications. For instance, Dijk et al. employed model-derived sleep and wake thresholds to examine the relationship between sleep timing and subjective sleepiness, as assessed by the Karolinska Sleepiness Scale (KSS), demonstrating that increased evening sleepiness was associated with earlier bedtimes and longer sleep duration [10]. Building on these insights, personalized light-based interventions have been developed to optimize sleep timing in aging populations using mathematical models [11]. The Homeostatic-Circadian-Light (HCL) model, which incorporates both sleep and wake thresholds, has further improved predictions of sleep timing and duration [12]. More recently, Hong et al. introduced the concept of circadian sleep sufficiency (CSS)–a metric integrating circadian-dependent thresholds–to address excessive daytime sleepiness in shift workers [13, 14]. Collectively, these advances underscore the importance of biologically meaningful thresholds in sleep-wake models and their translational potential for improving sleep health across diverse populations.

Despite these advances, a notable limitation of the PR model is its omission of the excitatory circadian projection to locus coeruleus (LC) mediated by orexin (Orx) neurons [15]. In the original formulation, circadian modulation of the sleep-wake switch acts solely via input to the VLPO [3, 16]. However, accumulating evidence indicates that SCN projections to Orx neurons play a critical role in stabilizing behavioral state transitions. Loss of Orx signaling–whether in knockout models or in individuals with narcolepsy–is associated with lowered sleep-wake thresholds and increased fragmentation of sleep architecture [17, 18]. Moreover, robust circadian rhythms in sleep-wake behavior persist even after VLPO lesions [19], and circadian dynamics remain intact in the absence of Orx signaling [20, 21]. These findings suggest that both inhibitory (to VLPO) and excitatory (to LC) projections contribute importantly to circadian regulation of sleep and wakefulness [15]. Therefore, explicitly modeling the excitatory circadian influence on Orx neurons is essential for a more complete understanding of sleep-wake dynamics.

To address this gap, we present a new sleep-wake model that incorporates the circadian drive’s excitatory input to Orx neurons. Within this framework, we redefine and compute circadian-dependent sleep and wake thresholds, enabling a more comprehensive description of the mechanisms underlying state transitions. We further apply this framework to investigate the effects of sleep deprivation and to preliminarily explore strategies for optimizing sleep-wake patterns in shift workers. Specifically, our objectives are fourfold: (i) to identify mechanisms governing sleep-wake transitions via bifurcation analysis and synchronization; (ii) to characterize the interaction between circadian drive and homeostatic sleep pressure by computing dynamic sleep and wake thresholds as functions of circadian phase; (iii) to demonstrate the effects of sleep deprivation using threshold-based analysis; and (iv) to optimize model parameters for healthy sleep in the context of two sleep-deprivation scenarios. In Section II, we briefly describe the structure and physiological basis of the proposed model. In the results (Sec. III), we present the dynamical mechanism of circadian rhythms in the modified sleep-wake model, the underlying nonlinear interaction between circadian and homeostatic drives, the recovery dynamics from sleep deprivation and finally apply the dynamical mechanism to the sleep-wake dynamics in shift workers. Finally, Sec. IV discusses our contributions and the implications of our findings, as well as future applications.

## II. MODIFIED FLIP-FLOP SLEEP-WAKE MODEL

Here we present a modified sleep-wake model in Fig. 1 that builds on the physiologically grounded PR model [8]. We begin with a conceptual overview of the relevant neurophysiology and model architecture, followed by a mathematical formulation of the system dynamics.

**FIG. 1:**
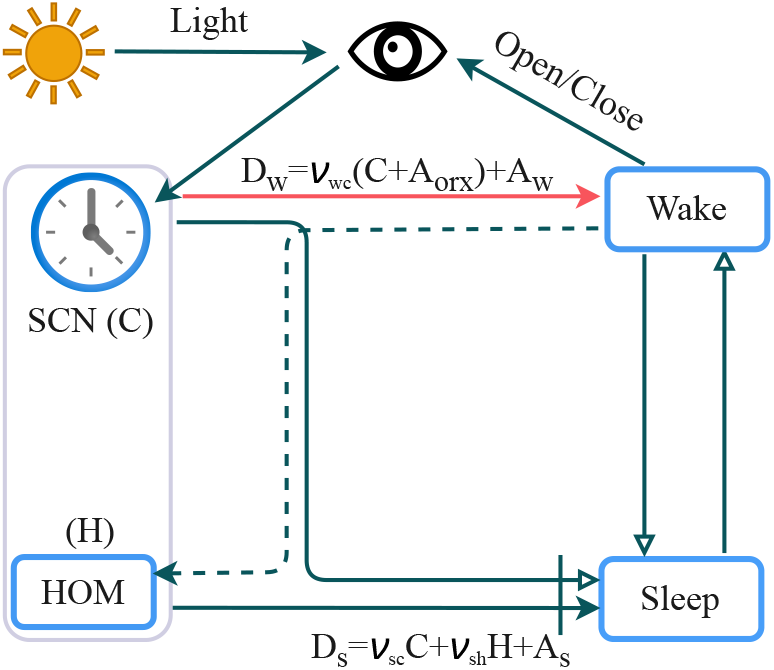
Schematic of the modified sleep-wake model based on the PR model [8]. The red pointer denotes the additional excitatory projection from the circadian drive to wake-promoting neurons introduced in our model.

The core model includes two reciprocally inhibiting neural populations: sleep-promoting and wake-promoting populations [22, 23]. These populations are regulated by two major processes: the circadian drive and the homeostatic sleep drive. The circadian drive, or Process C, originates from the SCN, which aligns internal timekeeping with the light-dark cycle via photic input [24]. The SCN influences sleep-wake regulation through two primary pathways: (1) It inhibits sleep-promoting neurons in the VLPO via projections through the dorsomedial hypothalamus (DMH) [3]; (2) It excites wake-active neurons, including Orx neurons and LC, via direct or indirect excitatory projections [17, 18]. Our model explicitly incorporates this latter excitatory circadian input to the wake-active population, which was absent in the original PR model.

The homeostatic drive reflects sleep need and is governed by the buildup and dissipation of sleep pressure [25, 26], often indexed by slow-wave activity (SWA). It increases during wakefulness–associated with the accumulation of adenosine–and decreases during sleep [27, 28]. Adenosine promotes sleep via A1 and A2A receptors; for instance, infusion of A2A agonists in certain brain regions induces sleep [29]. In this model, when adenosine (modeled as homeostatic pressure) exceeds a circadian-dependent sleep threshold, sleep-promoting neurons are activated. Conversely, when adenosine falls below the wake threshold, wake-promoting neurons regain dominance.

Figure 1 summarizes the model architecture. The dynamics of the wake population (*w*), sleep population (*s*), and homeostatic pressure (*H*) are governed by the following equations:

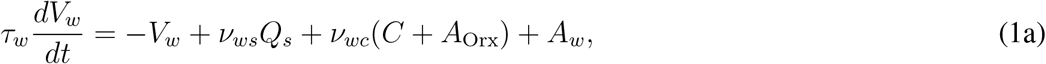

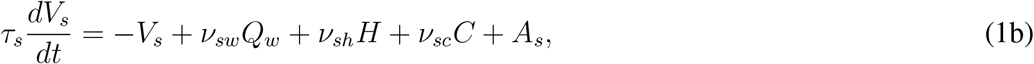

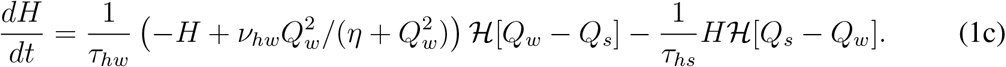

Here *V*_*a*_ (*a* = *w, s*) denotes the mean cell-body potential of Wake-active (*w*) and sleep-active (*s*) neurons. The firing rate *Q*_*a*_ is given by a sigmoid function:

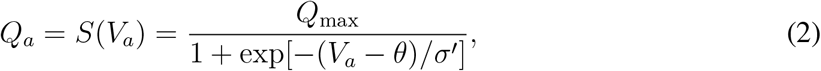

where *Q*_max_ is the maximum firing rate, *θ* being the mean firing threshold relative to resting, and 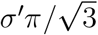 denoting the deviation [8]. The time constants *τ*_*a*_ (*a* = *w, s*) represent the characteristic neuromodulatory decay times. Interaction strengths population *b* to population *a* are denoted by *v*_*ab*_ for *a, b* = *w, s* (with *a* ≠ *b*) and are set as negative to capture mutual inhibitions between wake-active and sleep-active neurons.

For simplicity, we exclude light influence by modeling the circadian drive as a simple sinusoidal function, *C*(*t*) = sin(*ω*_*c*_*t*) with 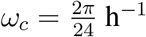, following previous studies on circadian rhythms [30, 31]. The amplitude is normalized to 1, as actual amplitudes can be incorporated into the interaction coefficients. The homeostatic drive (*H*), measured in percent SWA power [27, 28], increases (with *v*_*hw*_ *>* 0) during wake and decreases during sleep, mediated by adenosine accumulation. The Heaviside function ℋ[*x*] governs state dependency, being 0 for *x <* 0 and 1 for *x >* 0, where ℋ[*Q*_*w*_ − *Q*_*s*_] indicates the transition to wake. The parameter *v*_*sh*_ *>* 0 quantifies the excitatory influence of the homeostatic process on sleep-active neurons [8, 32]. External inputs *A*_*a*_ for *a* = *w*, Orx, *s* from other brain regions are set as constants. *A*_Orx_ represents constant input from the orexin groups, while *A*_*w*_ indicates constant input from the acetylcholine groups. The dynamics of homeostatic pressure are governed by two time constants: *τ*_*hw*_ for somnogen accumulation and *τ*_*hs*_ for its clearance [33]. A nonlinear function models the dependence of somnogen generation on *Q*_*w*_ with the parameters *η* and *v*_*hw*_. Nominal parameter values are presented in Tab. I.

**TABLE I:** Nominal parameter values.

| Parameter | Value | Unit | Parameter | Value | Unit |
| --- | --- | --- | --- | --- | --- |
| $Q_{\max}$ | 100 | $\text{s}^{-1}$ | $\nu_{ws}$ | -2.0 | $\text{mV s}$ |
| $\theta$ | 10 | $\text{mV}$ | $\nu_{sw}$ | -2.0 | $\text{mV s}$ |
| $\sigma'$ | 3 | $\text{mV}$ | $\tau_w$ | 10 | $\text{s}$ |
| $A_w$ | 1.4 | $\text{mV}$ | $\tau_s$ | 10 | $\text{s}$ |
| $A_{\text{Orx}}$ | 1.0 | $\text{mV}$ | | | |
| $A_s$ | 0 | $\text{mV}$ | $\nu_{sc}$ | -0.3 | $\text{mV}$ |
| $\eta$ | 2.3 | $\text{s}^{-2}$ | $\tau_{hw}$ | 45 | $\text{h}$ |
| $\nu_{hw}$ | 20 | $\text{nM}$ | $\tau_{hs}$ | 4.8 | $\text{h}$ |
| $\nu_{wc}$ | $0 \sim 3$ | $\text{mV}$ | $\nu_{sh}$ | $0 \sim 3$ | $\text{mV nM}^{-1}$ |

Compared to the original PR model, the key modification introduced here is the addition of an excitatory circadian input to the wake-promoting population (parameterized by *v*_*wc*_), representing SCN-driven excitation of Orx neurons. We also refine the homeostatic pressure dynamics to include a state-dependent, nonlinear accumulation term.

This model enables the systematic investigation of how circadian and homeostatic interactions shape sleep-wake transitions. In particular, we explore how variations in *v*_*wc*_ and *v*_*sh*_ influence the model’s behavior and thresholds. Other parameters are selected to preserve consistency with prior work on flip-flop sleep-wake switches [8, 34] and experimental observations under conditions such as sleep deprivation, fatigue, and circadian misalignment [35, 36].

## III. RESULTS

### A. Dynamical Mechanism of Circadian Rhythms

Although circadian rhythms are widely attributed to entrainment by external cues, the dynamical mechanisms that generate these rhythms within the sleep-wake system remain incompletely understood. In particular, how circadian and homeostatic drives interact to produce robust, cyclic transitions remains unsolved.

To explore this, we simulate the new model incorporating both circadian and homeostatic drives. Figure 2a displays the time series of wake- and sleep-promoting activity (*Q*_*w*_, *Q*_*s*_) for increasing homeostatic gain *v*_*sh*_, with circadian input *v*_*wc*_ fixed. Four distinct regimes emerge: 1) Low *v*_*sh*_: Prolonged wakefulness and lack of sleep pressure; 2) Moderate *v*_*sh*_: Short sleep episodes due to insufficient sleep pressure; 3) Nominal *v*_*sh*_: Canonical biphasic sleep-wake cycles (*~* 16h wake, *~* 8h sleep); 4) High *v*_*sh*_: Polyphasic sleep due to excessive sleep pressure. These results demonstrate that variations in *v*_*sh*_ alone reproduce the range of sleep dynamics seen across health and sleep disorders.

**FIG. 2:**
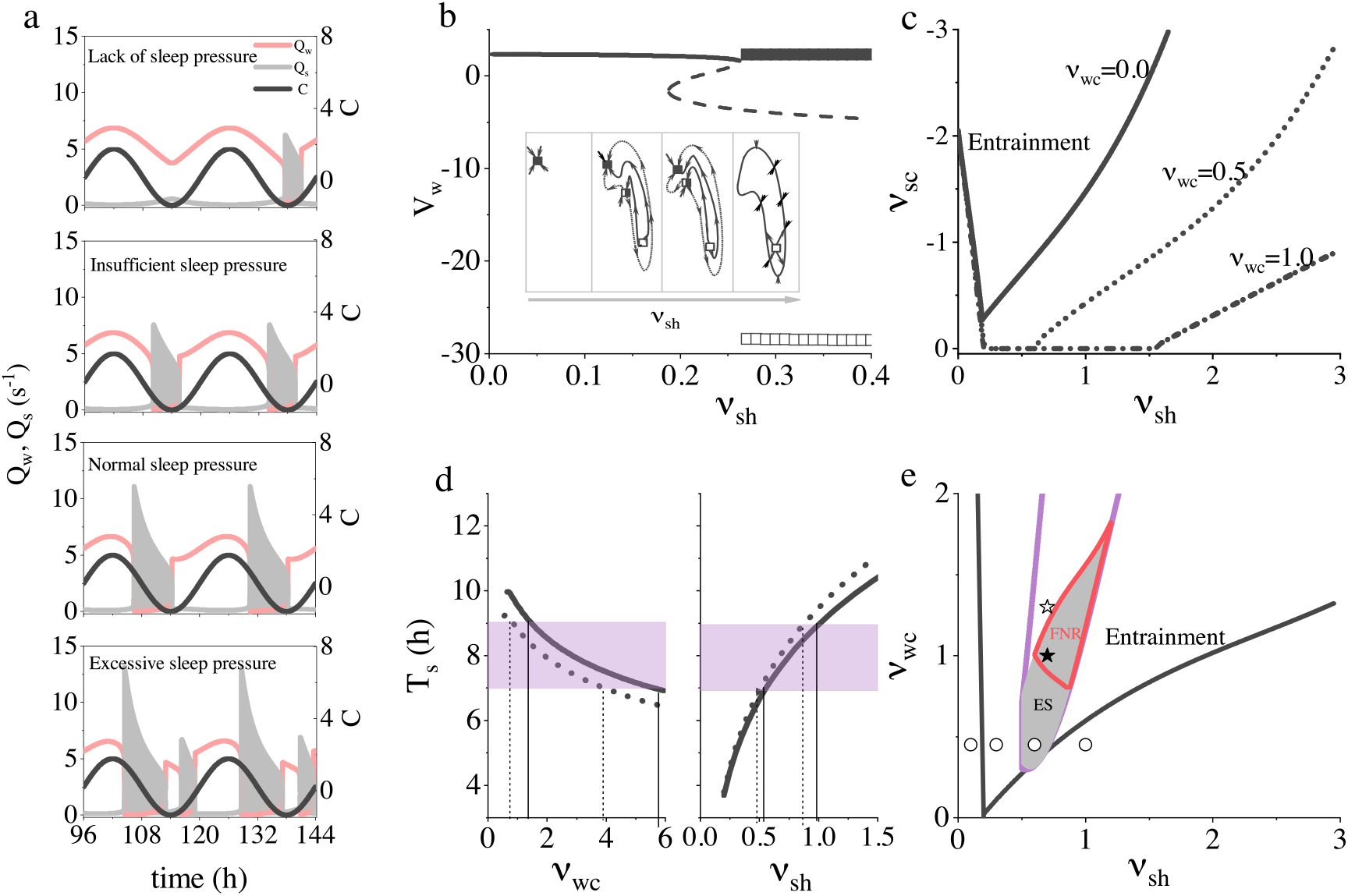
Dynamical mechanism of circadian rhythm and sleep-wake transitions. (a) Time series of wake (*Q*_*w*_) and sleep (*Q*_*s*_) activity for four values of homeostatic gain *v*_*sh*_ = 0.1, 0.3, 0.6, 1.0 with circadian input fixed at *v*_*wc*_ = 0.45 (denoted as open circles in (e)). Increasing *v*_*sh*_ leads to transitions from prolonged wakefulness to polyphasic sleep patterns. (b) Bifurcation diagram showing steady-state and oscillatory values of *V*_*w*_ versus *v*_*sh*_, with the circadian drive removed. An SNIC bifurcation gives rise to a limit cycle, marking the onset of rhythmic sleep-wake behavior (maxima: filled; minima: open squares), where the stable wake state (solid line) transitions become unstable (dashed line). Inset: phase space schematics showing the evolution of fixed points (filled: stable; half-filled: saddle with one + and two *−* eigenvalues; open squares: saddle with two + and one *−* eigenvalues; dashed line: the manifold connecting different states). (c) Entrainment regions in the (*v*_*sh*_, *v*_*sc*_) plane for various *v*_*wc*_. Entrainment (Arnold tongue) expands with stronger circadian input. (d) Sleep duration *T*_*s*_ against *v*_*wc*_ (left: dotted line, *v*_*sh*_ = 0.9; solid line, *v*_*sh*_ = 1.1) and *v*_*sh*_ (right: dotted line, *v*_*sh*_ = 1.2; solid line, *v*_*sh*_ = 0.8). (e) Phase diagram in the (*v*_*sh*_, *v*_*wc*_) plane. Black line: entrainment threshold; Purple-bordered area: normal sleep duration (7 *~* 9 h); Gray region: first type of sleep deprivation (no direct recovery); Red-bordered area: second type (partial recovery). ES: immediate sleep after deprivation; FNR: 10–30% first-night recovery. Hollow and solid stars: two deprivation cases in Fig. 4c.

To further dissect the underlying transitions, we perform a bifurcation analysis. By setting the circadian drive to zero (*v*_*sc*_ = *v*_*wc*_ = 0), we isolate homeostatic contributions. As shown in Fig. 2b, increasing *v*_*sh*_ shifts the system from a stable fixed point to oscillatory behavior via a saddle-node on invariant circle (SNIC) bifurcation [37]. The stable node, saddle, and unstable node coalesce, forming a heteroclinic loop and giving rise to a limit cycle (Fig. 2b, inset; more details in [37]). This SNIC mechanism offers a biologically grounded explanation for the emergence of rhythmic sleep-wake cycles from tonic wakefulness.

We then examine the system’s ability to entrain to a 24 h light-dark cycle, by analyzing the roles of the two circadian drive gains *v*_*sc*_ and *v*_*wc*_. Entrainment to the external cycle leads to the emergence of a robust 24 h sleep-wake rhythm, reflecting synchronization between the internal oscillator and the circadian input. This synchronization can be quantitatively characterized by the Arnold tongue in the (*v*_*sh*_, *v*_*sc*_) plane (Fig. 2c), which delineates the region where the intrinsic oscillator–arising via an SNIC bifurcation driven by homeostatic pressure–entrains to the external cue. In the absence of input to the wake-promoting population (*v*_*wc*_ = 0), a sufficiently large *v*_*sc*_ is required for entrainment, consistent with the original PR model. However, increasing 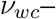 which amplifies circadian influence on wake-active neurons–expands the entrainment region and reduces the critical value of *v*_*sc*_, even allowing for successful synchronization when *v*_*sc*_ = 0. This indicates that the two circadian pathways act synergistically to stabilize the 24 h rhythm: stronger circadian input to either the sleep- or wake-promoting populations can compensate for weaker input to the other. These findings underscore the importance of coordinated circadian modulation of both neuronal populations in supporting the robust and flexible entrainment to environmental light-dark cycles, consistent with experimental observations described in Sec. I.

To this end, we focus on how the coupling between homeostatic sleep pressure and the circadian drive–mediated via orexin (Orx)–regulates sleep duration (Figs. 2d). The results show that increasing *v*_*sh*_ leads to longer sleep episodes, as a stronger homeostatic drive sustains activation of sleep-promoting neurons (Fig. 2d, left). Conversely, higher *v*_*wc*_ consolidates wakefulness by enhancing circadian input to wake-active populations, thereby reducing sleep duration (Fig. 2d, right). These findings delineate the parameter regimes that yield physiologically normal sleep durations (7 *~* 9 h) (purple shaded in Fig. 2d, and purple bordered area in Fig. 2e). Together, this analysis provides a comprehensive dynamical framework for understanding how homeostatic and circadian processes–and their coupling via orexinergic signaling–jointly orchestrate the timing and duration of sleep-wake cycles.

### B. Interaction between Circadian and Homeostatic Drives

While the circadian regulation of sleep-wake cycles is well understood, the precise interaction between circadian and homeostatic drives in governing state transitions remains unclear, motivating our analysis of their interplay within the sleep-wake regulatory network. Given the slow dynamics of homeostatic and circadian processes (*τ*_*hw*_, *τ*_*hs*_, *ω*^*−*1^ *≫ τ*_*w*_, *τ*_*s*_), we define two effective drive terms: *D*_*w*_ = *v*_*wc*_(*C* + *A*_Orx_) + *A*_*w*_ and *D*_*s*_ = *v*_*sc*_*C* + *v*_*sh*_*H* + *A*_*s*_, representing the net drives to wake-active and sleep-active neurons, respectively. Substituting into Eqs. 1a and 1b, the system reduces to:

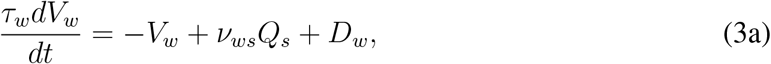

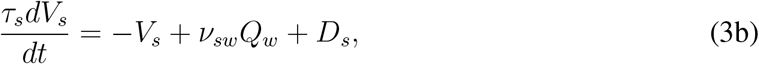

with steady-state values 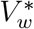 and 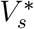 satisfying:

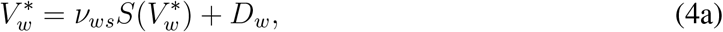

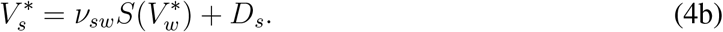

This leads to the implicit equation:

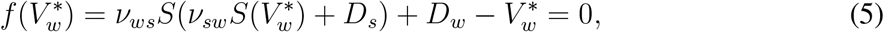

with an analogous expression for 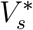.

The critical conditions for transitions between wake and sleep states are determined by locating saddle-node bifurcations, requiring that 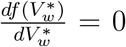. Solving this yields critical values of *D*_*s*_ as a function of 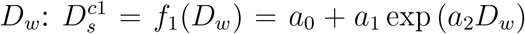 and 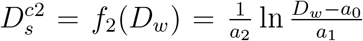, with *a*_0_ = *−*3.83742, *a*_1_ = 3.37745, *a*_2_ = 0.39306. These define the thresholds for transitions from wake to sleep and vice versa. Notably, when *v*_*ws*_ = *v*_*sw*_, *f*_2_ is the inverse of *f*_1_, reflecting symmetry in the system.

From these expressions, we derive explicit forms for the circadian-modulated thresholds of homeostatic sleep pressure *H*:

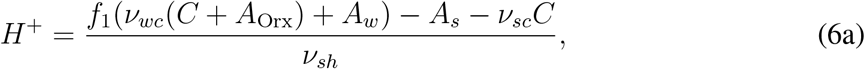

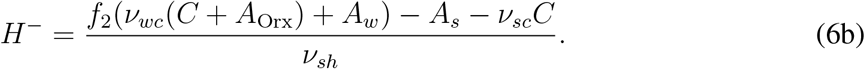

Here, *H*^+^ and *H*^−^ denote the thresholds for sleep onset and awakening, respectively. When *v*_*wc*_ = 0, these reduce to the thresholds of the original PR model [8, 13]. For *v*_*wc*_ *>* 0, the thresholds vary nonlinearly with circadian input, illustrating how circadian modulation dynamically shapes sleep-wake transition boundaries.

Figure 3 illustrates this interaction by plotting homeostatic sleep pressure *H* (blue line) along with the analytically derived circadian thresholds (Eq. 6). Sleep occurs when *H* exceeds *H*^+^ (gray shaded region), and wakefulness resumes when *H* drops below *H*^−^ (pink shaded region). These threshold crossings govern state transitions and capture the essential cyclical structure of the sleep-wake cycle. Comparing the three panels in Fig. 3, increasing *v*_*wc*_ elevates the circadian thresholds, delaying sleep onset and promoting longer wake periods while reducing total sleep duration. These results underscore the nonlinear, synergistic interaction between circadian and homeostatic drives. Interestingly, despite variation in individual parameter values, similar sleep-wake patterns can arise from distinct parameter sets–a phenomenon known as *parameter degeneracy*. This suggests that multiple combinations of physiological mechanisms may give rise to comparable behavioral outcomes. However, this degeneracy poses challenges for identifying biologically accurate parameter regimes. Resolving this ambiguity will require future work incorporating empirical constraints and data-driven parameter estimation.

**FIG. 3:**
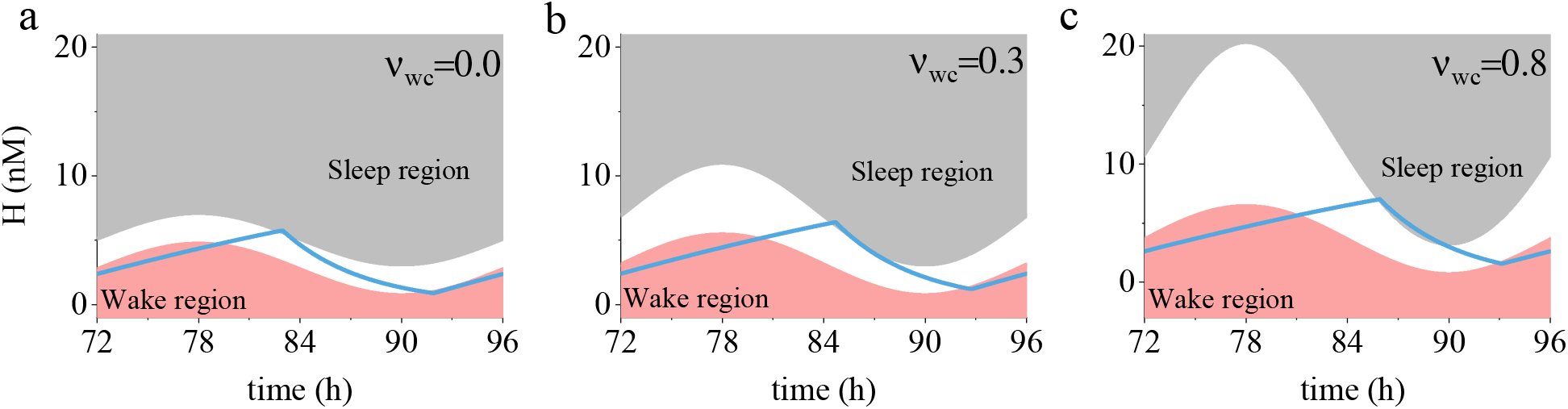
Interaction between homeostatic sleep pressure (blue line) and circadian drive during the sleep-wake cycle is illustrated. Gray border: circadian sleep threshold; Pink border: circadian wake threshold. From left to right, increasing circadian gain *v*_*wc*_ to wake-promoting neurons.

### C. Sleep Deprivation

To further probe the physiological implications of circadian-homeostatic interaction, we investigate system behavior under sleep deprivation. Sleep deprivation presents a well-established paradigm for testing the resilience, and adaptability of sleep-wake regulation and understanding its underlying dynamical mechanisms for linking model predictions to empirical phenomena. To accurately model subjective fatigue and evaluate the system’s recovery capacity following sleep loss, we examine whether the model meets two key experimental criteria: 1) Immediate sleep onset following prolonged deprivation; 2) Recovery sleep compensating for approximately 10–30% of the lost sleep, as consistent with empirical observations [8].

#### 1. Analytical predictions

For the first criterion, we fix *Q*_*w*_ and *Q*_*s*_ to simulate enforced wakefulness, maintaining a sustained wake state from *t*_0_ to *t*_0_ + 80 h, during which the circadian drive *C* reaches its peak. The homeostatic pressure *H* evolves according to:

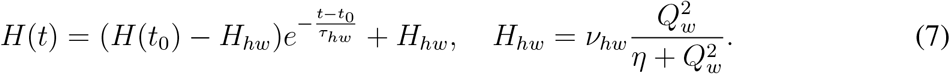

Then the sleep pressure at deprivation end can be given as:

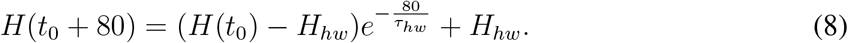

Sleep onset requires *H*(*t*_0_ + 80) to exceed the circadian-modulated threshold *H*^+^:

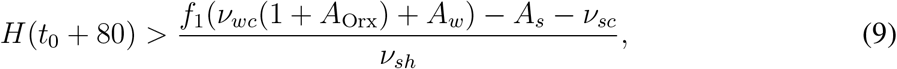

which leads to the critical condition for *v*_*sh*_:

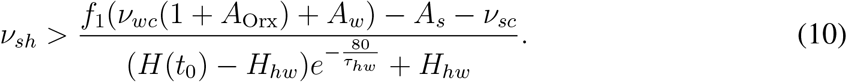

For the second criterion, we model recovery from a 60 h deprivation starting at normal sleep onset 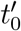. The homeostatic pressure at 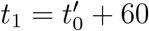 can be given as:

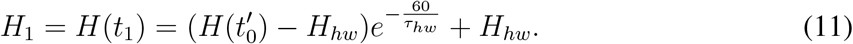

Recovery sleep duration *T*_first_ can be determined by the condition *H*(*t*_1_ + *T*_first_) = *H*^−^, which yields:

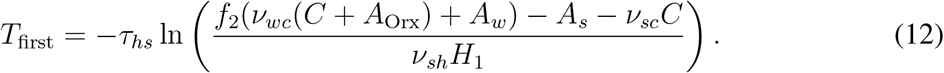

This predicts *T*_first_ to exceed the baseline sleep duration by 10–30%, consistent with experimental data [8].

These theoretical predictions provide explicit dynamical and physiological constraints for sleep-wake transitions during sleep deprivation and recovery, allowing for a mechanistic interpretation of circadian-homeostatic interaction under substantial challenge.

#### 2. Numerical results

We tested these predictions via simulation. For the first scenario under 80h deprivation (Fig. 4a), low *v*_*wc*_ enables immediate sleep at the end of deprivation (upper panel), while high *v*_*wc*_ prevents sleep onset (lower panel), due to elevated circadian sleep thresholds (Fig. 4b). These effects are recapitulated in the (*D*_*w*_, *D*_*s*_) plane (Fig. 4c): the low-*v*_*wc*_ trajectory crosses the sleep boundary *D*_*s*_ = *f*_1_(*D*_*w*_), while the high-*v*_*wc*_ trajectory remains outside the sleep domain. These results corroborate the theoretical predictions in Sec. III C 1. For the second scenario, the 60h deprivation case (Fig. 4d), the model exhibits extended recovery sleep, and the ratio *R* = *T*_first_*/T*_baseline_ depends sensitively on model parameters (Fig. 4e). Higher *v*_*wc*_ reduces *R* by promoting early awakening, while larger *v*_*sh*_ enhances recovery sleep and maintains *R >* 30%.

**FIG. 4:**
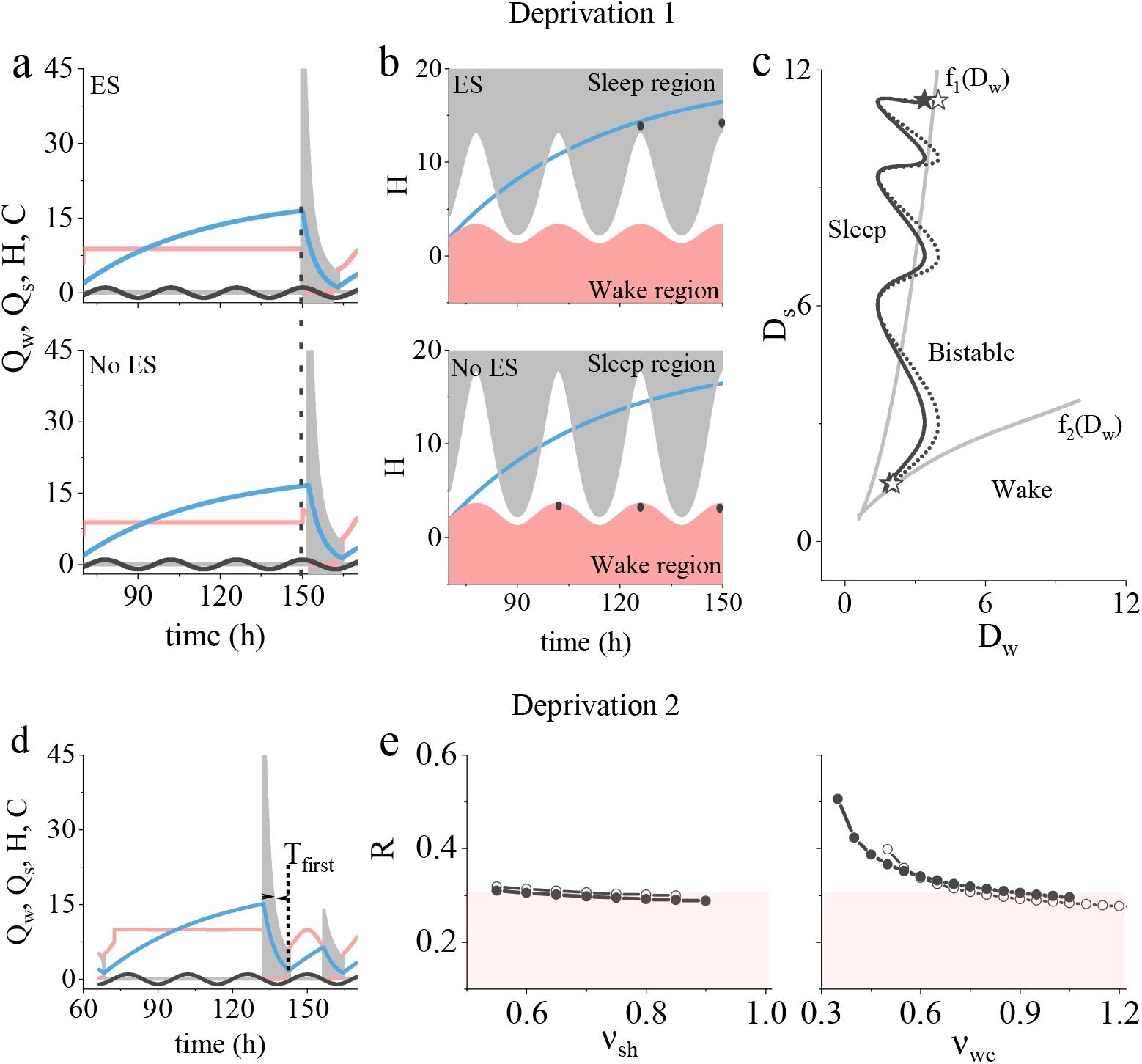
Model behaviors under sleep deprivation. (a) Model response to 80 h sleep deprivation for two values of *v*_*wc*_: *v*_*wc*_ = 1.0 (top) and 1.3 (bottom) with *v*_*sh*_ = 0.7. Parameters are denoted by hollow and solid stars in Fig. 2e. Dashed line indicates deprivation end, aligned with circadian peak. (b) Time course of homeostatic pressure (black), circadian sleep threshold (gray), and wake threshold (pink). “ES” and “No ES” indicate whether immediate sleep occurs post-deprivation. (c) Dynamical phase diagram in (*D*_*w*_, *D*_*s*_) plane. Colored lines show trajectories during the deprivation period; markers indicate the corresponding parameter sets in Fig. 2e. (d) Model output following 60h deprivation initiated at habitual sleep onset, showing first-night recovery sleep at *v*_*wc*_ = 1.0, *v*_*sh*_ = 0.7. (e) Ratio *R* of recovery sleep duration (*T*_first_) to baseline: (e_1_) *R* as a function of *v*_*sh*_ with *v*_*wc*_ = 0.8 (circles) and 0.9 (dots); (e_2_) *R* as a function of *v*_*wc*_ with *v*_*sh*_ = 0.6 (circles) and 0.8 (dots).

These findings highlight how the dynamic interplay between homeostatic and circadian drives constrains the system’s response to sleep deprivation and and its ability to recover from extreme sleep loss. The proposed framework quantitatively captures the trade-offs and compensatory mechanisms shaping sleep rebound and reveals key parameter regimes that support physiologically realistic sleep dynamics.

### D. Sleep-Wake Dynamics in Shift Workers

Shift work, now undertaken by approximately 20% of the workforce [38], often imposes schedules misaligned with natural circadian rhythms, increasing the risk of sleep disturbances, fatigue, and accidents [39, 40]. To examine how such misalignment affects sleep-wake regulation, we simulate the model’s response to a 120-day periodic light exposure protocol commonly used in experimental studies of shift workers. This setup enables us to evaluate how the interaction between circadian and homeostatic processes governs sleep-wake transitions and to assess the system’s adaptability to disrupted schedules, offering insights into mechanisms underlying shift work-related risks.

Under a standard 16:8 light-dark (LD) cycle (250 lux light, 0 lux dark), the model establishes a stable baseline pattern–nighttime sleep and daytime wakefulness–entrained by sinusoidal circadian drive (*ω*_*c*_ = 2*π/*24 h^*−*1^), thus aligning the internal clock with the external environment.

However, under irregular lighting conditions that deviate from the natural LD cycle (Fig. 5, top), this alignment breaks down. The resulting desynchronization between circadian oscillators and external light cues manifests as pronounced deviations in the firing rates of wake- and sleep-active neurons (Fig. 5, middle). These irregular sleep-wake patterns are further clarified by analyzing homeostatic sleep pressure in relation to circadian-dependent sleep and wake thresholds (Fig. 5). Due to misaligned schedules, sleep pressure (*H*) exhibits aperiodic fluctuations and often fails to exceed the circadian wake threshold needed to initiate wakefulness, leading to truncated or insufficient sleep episodes. This misalignment disrupts the coordination between circadian and homeostatic drives, ultimately compromising the timing, consolidation, and quality of sleep.

**FIG. 5:**
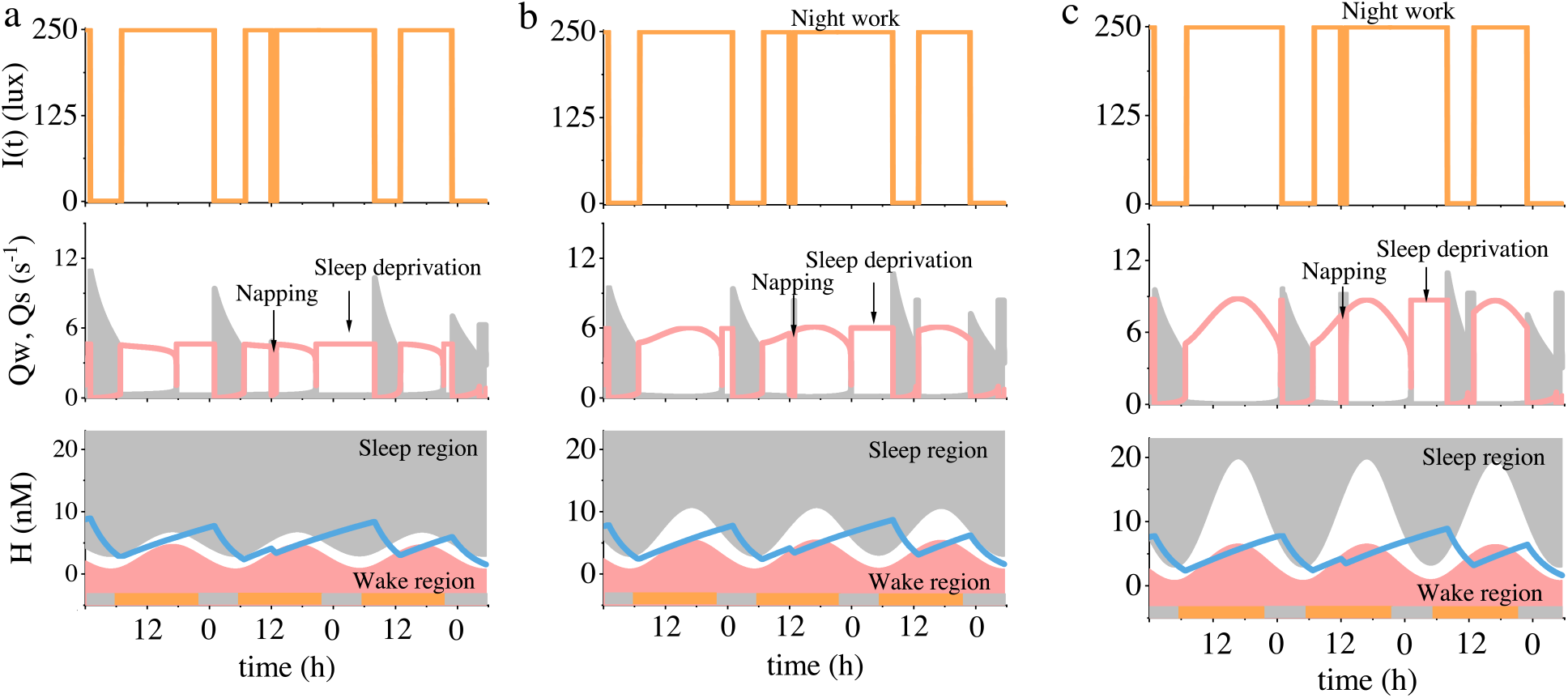
Model simulation of shift workers’ sleep-wake dynamics after 120 days periodic light exposure. Top: imposed light-dark cycle; Middle: neuronal firing rates *Q*_*w*_ (pink) and *Q*_*s*_ (gray); Bottom: homeostatic sleep pressure *H* (black), circadian sleep threshold (pink), and wake threshold (blue). From left to right: *v*_*wc*_ = 0.0, 0.3, 0.8 with *v*_*sh*_ = 0.4, *v*_*sc*_ = −0.8.

By comparing the three subplots (Fig. 5, middle), we observe that the duration of sleep deprivation decreases with increasing *v*_*wc*_–evidenced by the shortening of horizontal wake periods across the panels. This trend is further corroborated by the corresponding interaction diagrams in Fig. 5, bottom. Although the homeostatic sleep pressure trajectories remain consistent across all three conditions (due to identical *v*_*sh*_ values), the elevation of the circadian sleep threshold with higher *v*_*wc*_ delays the onset of sleep. As a result, less homeostatic pressure is required to trigger sleep onset, effectively reducing the length of sleep deprivation. Interpreting the deprivation duration as a proxy for excessive daytime sleepiness (EDS), our results suggest that stronger circadian input–especially through orexin-mediated wake drive–can mitigate EDS and enhance sleep quality. While these findings are qualitatively consistent with prior work [41], our model introduces a key refinement: the use of dynamic, circadian-dependent sleep and wake thresholds. This necessitates a reevaluation of the CSS metric originally introduced in Ref. [13], which was based on fixed thresholds.

## IV. DISCUSSIONS AND CONCLUSIONS

In this work, we developed a modified PR model incorporating *circadian-dependent sleep and wake thresholds* to investigate the dynamical interactions between the circadian rhythm (Process C) and homeostatic sleep pressure (Process S). Through a combination of bifurcation analysis, theoretical analysis, and numerical simulations, we examined how this coupling gives rise to normal, deprived, and shift-workers’ irregular sleep patterns, offering a more physiologically grounded understanding of sleep-wake regulation.

Our main findings are as follows:

### 1) Dynamical Mechanism of Circadian-Homeostatic Coupling

In the absence of circadian input, bifurcation analysis shows that increasing the homeostatic parameter *v*_*sh*_ drives transitions from sustained wakefulness to monophasic and polyphasic sleep, eventually yielding cyclic sleep-wake oscillations via an SNIC bifurcation. The addition of a circadian drive (either to VLPO or to LC) entrains the cycle to a 24h rhythm, with *Arnold tongue* structures delineating regions of synchronization. Robust, physiologically realistic rhythms require sufficiently strong circadian and homeostatic contributions; disruption in either component leads to irregular, fragmented sleep.

### 2) Interaction of Circadian and Homeostatic Drives

By analytically deriving dynamic, circadian-modulated sleep and wake thresholds based on net drives, our model captures the nonlinear interaction between homeostatic pressure and circadian signals. Notably, stronger circadian input (parameterized by *v*_*wc*_) elevates the sleep threshold, consolidating wakefulness and reducing sleep duration. Despite this, parameter degeneracy–where different combinations of *v*_*sh*_ and *v*_*wc*_ yield similar sleep patterns–highlights the need for experimental constraints to uniquely determine biologically plausible parameter sets.

### 3) Sleep Deprivation and Recovery Dynamics

Our model replicates hallmark features of sleep deprivation: immediate sleep onset following extended wakefulness and partial rebound sleep (10-30%) during subsequent recovery, consistent with experimental data. Elevated circadian drive delays post-deprivation sleep onset and shortens recovery, while increased homeostatic drive prolongs rebound sleep. These findings provide mechanistic insight into how the circadian and homeostatic systems respond to extreme sleep loss and recovery.

### 4) Shift Work and Circadian Misalignment

Simulating shift work under irregular lighting schedules reveals that higher circadian strength (*v*_*wc*_) shortens periods of sleep deprivation but induces phase shifts and sleep fragmentation when misaligned with external light cues. These dynamics are driven by the newly introduced *circadian-modulated thresholds*, which adaptively shift based on light-dependent circadian input. This framework surpasses classical models by accounting for delayed sleep onset, early awakening, and irregular sleep patterns under real-world disruptions like shift work or jet lag.

The introduction of dynamic, circadian-modulated sleep and wake thresholds marks a substantial advancement in computational models of sleep-wake regulation. Traditional models [1, 8] often relied on static or heuristically defined thresholds, limiting their physiological realism. By embedding circadian modulation directly into the criteria for sleep onset and awakening, our model captures a wider range of biologically relevant phenomena and allows for a more precise quantification of circadian misalignment and its consequences. This refinement enables the systematic exploration of how endogenous rhythms interact with environmental and behavioral disruptions, and supports investigations into conditions such as sleep fragmentation [42], delayed sleep onset [43], early awakening [44], excessive daytime sleepiness [45], and insomnia [46]. It also provides a concrete basis for recalculating metrics like the CCS [13] using our updated threshold definitions, thus potentially improving the accuracy of sleep assessments under shift work conditions.

Our findings carry important implications for both basic and applied sleep science. Misalignment between circadian and homeostatic processes–whether due to shift work, jet lag, or genetic variation–can contribute to a range of sleep disorders. Our results suggest that interventions targeting circadian thresholds or their alignment with homeostatic signals may offer promising strategies for improving sleep health [47].

While our model offers valuable insights, several directions remain for future work: 1) Incorporating environmental and lifestyle factors: Integrating influences such as light exposure, diet, and exercise could enhance the model’s ecological validity and support personalized, non-pharmacological interventions based on the circadian-homeostatic interaction. 2) Modeling individual variability Accounting for genetic or chronotype-related differences in circadian dynamics and sleep behavior may improve the model’s applicability to diverse populations. 3) Experimental validation: Using physiological recordings and behavioral data to further constrain parameter ranges and validate model predictions will be critical for translation to clinical and occupational settings.

## V. MATERIALS AND METHODS

In this model, the circadian drive *C* (dimensionless) originates from the SCN and is synchronized to an approximately 24.15-hour cycle via retinal photoreceptor input [48]. The drive *C* influences the sleep-promoting neurons (e.g., in the VLPO) through inhibitory signals (*v*_*sc*_ *<* 0), while wake-promoting centers (e.g., Orx and LC) receive excitatory inputs (*v*_*wc*_ *>* 0) from the circadian system [3, 17, 18, 22]. We model the circadian drive as:

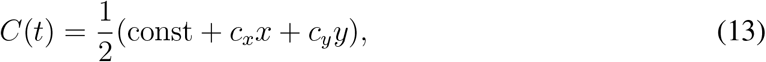

where *x* and *y* are the state variables of a forced, modified van der Pol oscillator [49]:

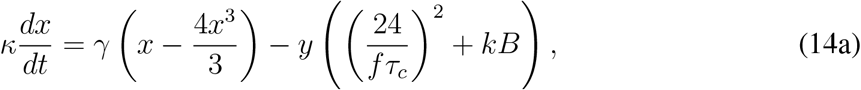

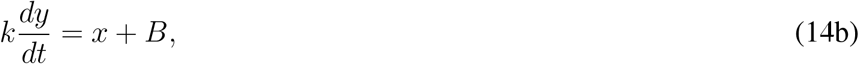

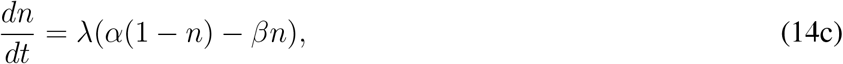

with *n* being the fraction of activated photoreceptors and *B* = *Gα*(1*−n*)(1*−bx*)(1*−by*) representing the drive from the photoreceptive pathway to the circadian pacemaker, where 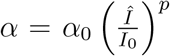 and the light exposure *Î*described as follows:

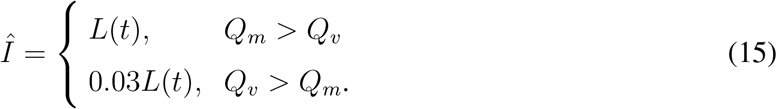

where *L*(*t*) is the ambient light level and wake is defined by *Q*_*m*_ > *Q*_*v*_, the data for light exposure comes from Ref. [13]. For the circadian drive: const = 0.0, *c*_*x*_ = −0.16, *c*_*y*_ = 0.8, *τ*_*c*_ = 24.09, *κ* = (12*/π*), *i*_0_ = 9500, *p* = 0.6, *γ* = 0.23, *k* = 0.55, *β* = 0.013, *f* = 0.99669, *α*_0_ = 0.16, *b* = 0.4, *G* = 19.9.

## Acknowledgments

This work was supported partially by the Natural Science Foundation of Zhejiang Province under Grant Nos. LY24A050003, the National Natural Science Foundation of China under Grant Nos. 12175242.

